# The structure of a *Plasmodium vivax* Tryptophan Rich Antigen suggests a lipid binding function for a pan-*Plasmodium* multi-gene family

**DOI:** 10.1101/2022.09.30.510049

**Authors:** Prasun Kundu, Deboki Naskar, Shannon McKie, Sheena Dass, Usheer Kanjee, Viola Introini, Marcelo U. Ferreira, Manoj Duraisingh, Janet Deane, Julian C. Rayner

## Abstract

Tryptophan Rich Antigens (TRAgs) are encoded by a multi-gene family in all *Plasmodium* species, significantly expanded in *P. vivax*, but their function is not currently known. We show that multiple *P. vivax* TRAgs are expressed on the merozoite surface and that one, PVP01_0000100 binds red blood cells with a strong preference for reticulocytes. Solving the structure of the C-terminal tryptophan rich domain that defines the TRAg family revealed a three-helical bundle that is conserved across *Plasmodium* and has homology with lipid-binding BAR domains involved in membrane remodelling. Biochemical assays confirmed that this domain has lipid binding activity with preference for sulfatide, a glycosphingolipid present in the outer leaflet of plasma membranes. Deletion of the putative orthologue in *P. knowlesi*, PKNH_1300500, impacts invasion in reticulocytes, suggesting a role for membrane remodelling during this essential process. Together, this work suggests a molecular function for the TRAg family for the first time.

## Introduction

*Plasmodium vivax* is the malaria parasite with the broadest geographic distribution, with nearly 3 billion people at risk of infection, primarily in Latin America and Asia[1]. It differs from *P. falciparum* in multiple aspects of its biology, most notably by the presence of the hypnozoite, a dormant liver stage can lead to recurrent relapses even in the absence of transmission, and the fact that sexual stage gametocytes form much more rapidly, meaning that transmission occurs more frequently and is harder to control. As a result, *P. vivax-*specific intervention strategies are a specific goal for the WHO Malaria Control Plan [2] However, progress in understanding *P. vivax* biology has been slowed in part by its absolute requirement for growth in immature red blood cells called reticulocytes, which has hampered development of *in vitro* culture systems.

The process by which *Plasmodium* parasites identify and invade human red blood cells has long been of interest for vaccine development, as the process is essential for parasite survival and therefore malaria pathology. In *P. falciparum*, the essential invasion protein PfRh5, which binds the erythrocyte receptor Basigin, is a high priority vaccine target[3, 4] and Phase IIa trials show that even early-stage PfRh5 vaccines can reduce parasite growth rates *in vivo*[5]. The phylogenetic distance between *P. falciparum* and *P. vivax* means that there is no clear identifiable orthologue of PfRh5 in *P. vivax*, and blood-stage vaccine attention has focussed primarily on Duffy Binding Protein, PvDBP, with early stage vaccine trials under way[6]. However, sequence and copy number variation in PvDBP between *P. vivax* isolates is well-known[7] raising the potential for vaccine escape and emphasising that other blood-stage targets also need to be investigated. Of particular interest are *P. vivax* proteins that specify reticulocytes for invasion. The recognition of reticulocytes depends, at least in part, on the interaction between *P. vivax* Reticulocyte Binding Protein 2b (PvRBP2b) and CD71/Transferrin Receptor[8], but the process of red blood cell invasion is a highly complex, multi-stage process[9], which likely involves additional proteins.

Potential candidate proteins include the Tryptophan rich antigen multi-gene family (TRAgs), which all share a 254-536 amino acid long Tryptophan-Threonine-rich plasmodium antigen C terminal (tryThrA_C) domain of unknown function (PFAM ID PF12319, OrthoMCL ID OG6_145873), usually at their C-terminus. First identified in the rodent model *Plasmodium yoelii*, TRAgs have been shown to generate highly protective antibodies in mice upon immunisation [10, 11]. All *Plasmodium* species possess some TRAgs, but they are particularly numerous in *P. vivax* and closely related species, where they are frequently found in clusters along chromosomes, indicative of expansion through gene duplication and diversification. The precise function of TRAgs in not known in any *Plasmodium* species, but several members of the *P. vivax* TRAg family have been reported to have red blood cell binding properties[12] and in some cases interacting partners have been reported[13, 14]. TRAgs also differ in expression between *P. vivax* infections in *Samiri* and *Aotus* monkeys, suggestive of a variable role, potentially as alternative ligands to PvDBP[15]. Members of the *P. vivax* TRAg family are highly immunoreactive, with a mean seropositivity rate of around 60% even in samples from low endemic regions [16].

To explore the function of *P. vivax* TRAgs in a systematic manner, we exploited a eukaryotic recombinant protein expression system that we have previously employed to identify the interaction between PfRh5 and Basigin[3] and to screen for *P. vivax* vaccine candidates [17, 18]. Antibodies raised against multiple *P. vivax* TRAgs suggested a merozoite surface location and revealed errors in genome annotation. Focusing on one family member, PVP01_0000100, we established that it bound to both erythrocytes and reticulocytes, with a preference for the later, via the C-terminal domain. Solving the crystal structure of this domain revealing an extended, curved three-helical bundle, with similarity to lipid binding BAR domains. Lipid and liposome binding assays confirmed that the PVP01_0000100 TRAg domain bound specifically to sulfatide, and deletion of the homologous gene in *P. knowlesi* resulted in a decrease in reticulocyte preference in invasion assays. This combination of biochemistry, structural biology and experimental genetics reveals a molecular function for *Plasmodium* TRAgs for the first time.

## Results

### Tryptophan Rich Antigens (TRAgs) are expanded in some *Plasmodium* species

To facilitate a systematic exploration of the TRAg family we used three different methods to identify TRAgs: searching PFAM and OrthoMCL databases with the relevant domain ID for their conserved Tryptophan/Threonine-rich domain (PF12319 and OG6_145873 respectively) and BLAST searching PlasmoDB using the C-terminal domain from PVP01_0000100, eliminating low confidence hits. The numbers of TRAgs identified differed depending on the method used (**Table 1**), as the databases all use different reference genomes and different domain definitions. However, it is clear that the TRAg family, while present in all *Plasmodium* species, has expanded significantly in the primate *Plasmodium* clade that includes four of the five major human *Plasmodium* species - *P. vivax, P. knowlesi, P. ovale* and *P. malariae*. Multiple different nomenclatures have been used for the TRAg family, which complicates analysis and comparison [16, 19]. To facilitate understanding we use the current *P. vivax* reference genome gene IDs throughout this work and provide a systematic summary of all nomenclature systems in **Supplementary Table 1** A neighbour joining tree based on full-length TRAg sequences from *P. vivax, P. knowlesi* and *P. cynomolgi* (**Figure 1A**) reveals that many genes have 1:1:1 orthologue in all three species, suggesting these TRAgs originated in a common ancestor before the three species diverged. There were however also examples of closely related genes in *P. vivax* and *P. cynomolgi* with no homologue present in *P. knowlesi*, suggesting they originated since the divergence of a common ancestor of *P. vivax* and *P. cynomolgi* from *P. knowlesi*, as well as *P. vivax* TRAgs with no clear 1:1 orthologue in either species, suggesting an origin since the split of *P. vivax* and *P. cynomologi*. This pattern suggests the process that led to the expansion of the TRAg family in the *Plasmodium* subgenus is still occurring, with new genes evolving as *Plasmodium* species diverge.

**Table 1:**
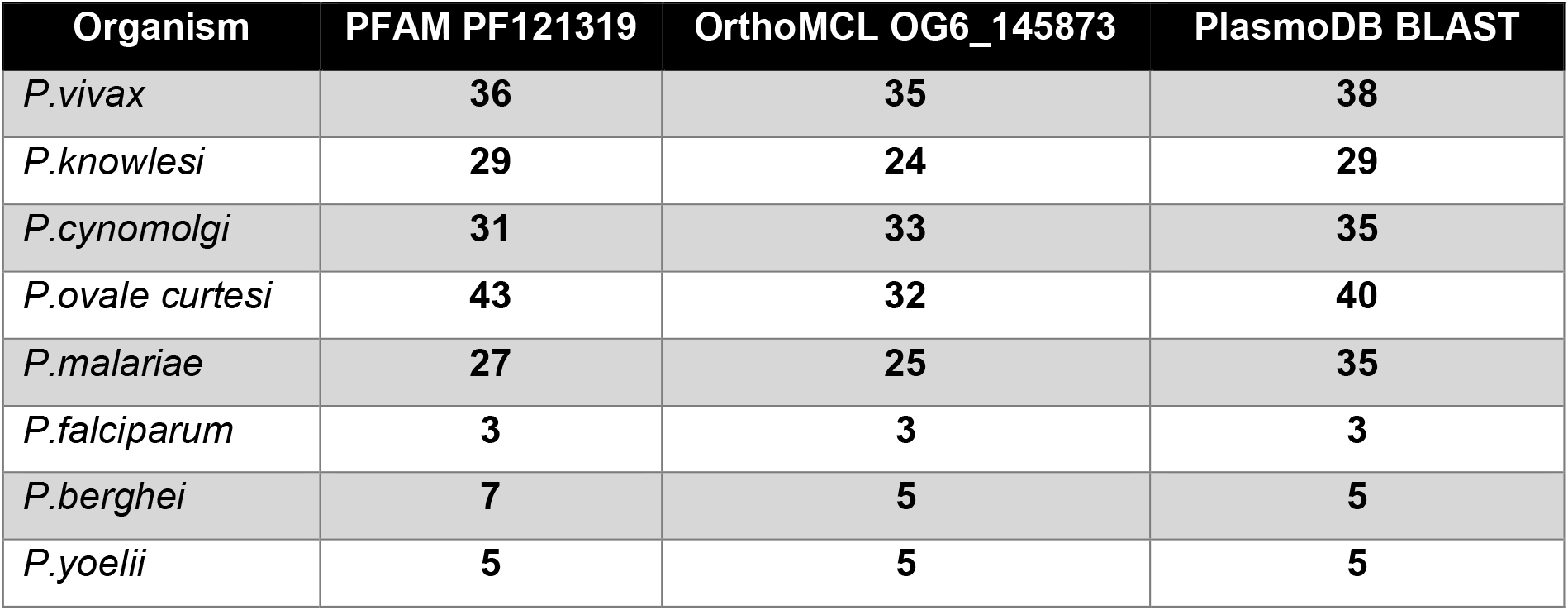
Expansion of the TRAgs in specific *Plasmodium* species.

**Figure 1:**
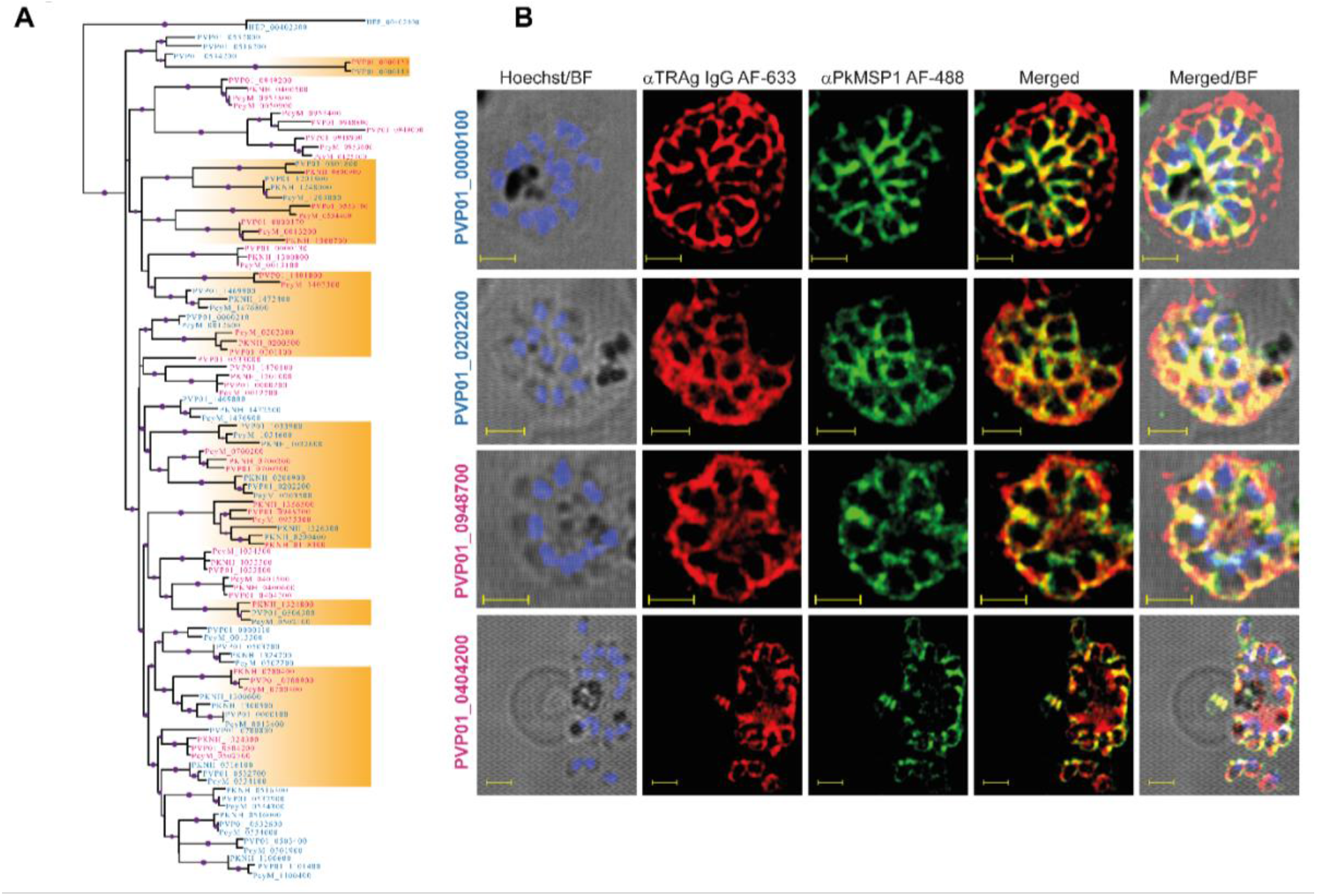
Multiple PvTRAGs localise to the merozoite surface, regardless of signal peptide prediction. **A) Phylogenetic tree of TRAgs**. Pv, Pk and Pc TRAgs (colour coded as Blue for -SP and magenta +SP) show their position on the phylogenetic tree. The phylogenetic tree was reconstructed using the maximum likelihood method with 100 bootstrap replicates. The scale for the bootstrap values is 0.25-1 represented with the size of the purple spherical symbol on each branch. The orange boxes highlight clusters where TRAgs with and without predicted SPs are grouped together. This suggests the SPs must have been gained or lost multiple times in the evolution of the family, which seems unlikely – issues with genome annotation, discussed in the text, seem more likely. **B) Immunolocalisation of TRAgs on *P. vivax ex vivo* isolates**. Rabbit polyclonal IgGs raised against four PvTRAgs (+/- SP) were used to study the localisation of the proteins in *P. vivax* isolates. A rat antibody against PkMSP_1-19_ was used as a merozoite surface marker for co-localisation studies. Alexa Fluor 633 goat-anti rabbit and Alexa Fluor 488 goat-anti rat were used as secondary antibodies for staining. Hoechst 33342 was used to stain the nucleus. The first column is an overlay of Hoechst with brightfield images, while the last column is an overlay of Alexa Fluor 633, Alexa Fluor 488, Hoechst and brightfield (BF). The highly colocalised fluorescence intensity of the green and red channel indicates the presence of the TRAgs on the surface of the individual Pv merozoites. The colocalised fluorescence intensities are quantified in **Supplementary Figure 2**. Scale bars are 2 microns.

### Some *P. vivax* TRAgs are localised on the merozoite surface irrespective of whether they are predicted to contain a signal peptide

In keeping with previous analysis [16] some of the *P. vivax* TRAgs we identified are predicted to have SPs in PlasmoDB, but some are not. However, all predictions in PlasmoDB are made using one SP prediction server, SignalP[20] Analysis of *P. vivax* TRAg sequences using alternative servers, Phobius[21] and PredSci[22], identified potential SPs in several TRAgs that are not identified by SignalP (**Supplementary Table 2**). Notably, all *P. vivax* TRAGs lacking a predicted SP instead contained a domain annotated as ‘N-terminal transmembrane’, which we hypothesised may actually function as an SP. To test this, we generated polyclonal antibodies in rabbits against four PvTRAgs, both with and without predicted SPs. Antibodies were raised against full-length recombinant proteins expressed in the HEK293E cell system that has been used extensively for *Plasmodium* protein expression [23], and purified using affinity chromatography. To test for cross-reactivity, we used the antisera to probe Western blots containing a panel of 12 purified recombinant *P. vivax* TRAgs. Since all the constructs have a Biolinker peptide and hexa-His tag in common, some level of background was expected and Pfs25, which includes the same tag, was used as background negative control. Only a very low-level of cross-reactivity was found for the anti-PVP01_0000100, PVP01_0532700 and PVP01_0202200 antisera, while anti-PVP01_0404200 had some level of cross-reactivity with one other TRAg, PVP01_0202200 (**Supplementary Figure 1**).

Immunofluorescence assays using *ex vivo P. vivax* parasites enriched for schizonts showed merozoite surface localisation for all antisera, whether they were raised against PvTRAgs without (PVP01_0000100 and PVP01_0202200) or with (PVP01_0948700 and PVP01_0404200) a predicted SP. Merozoite surface localisation was defined based on co-localisation with merozoite surface stain marker PkMSP_1-19_ **(Figure 1B)**, with the Pearson Correlation Coefficient (r) for colocalization averaging 0.8 (**Supplementary Figure 2A**). All four anti-PvTRAg antibodies also yielded predominantly merozoite surface location in *P. knowlesi*, again irrespective of whether the TRAgs were predicted to contain an SP or not (**Supplementary Figure 3)**. The r value for colocalization with PkMSP1 was lower in *P. knowlesi*, presumably because of lower cross-reactivity of anti-PvTRAg antibodies with their *P. knowlesi* orthologues (**Supplementary Figure 2B**). Together, these data support the hypothesis that the lack of predicted SP in many TRAgs is an artefact of automated genome annotation and potentially all members of the TRAg family enter the secretory pathway.

### PVP01_0000100 binds preferentially to human reticulocytes via its C-terminal domain

Given the location of multiple TRAgs on the merozoite surface (**Figure 1B** and refs [14, 24]) we hypothesised that they may bind directly to the surface of erythrocytes or reticulocytes and play a role in parasite invasion. To test this, we trialled expression of a library of 33 PvTRAgs using the HEK293E system where full length ectodomains are secreted into the culture supernatant, from which they can detected and purified using a C-terminal biotinylation and hexa-His tag. Expression of 27 PvTRAgs was confirmed by Western blot using HRP conjugated anti-His antibody **(Supplementary Figure 4)**. Sufficient yield for binding studies was obtained for 15 PvTRAgs. To increase avidity, affinity purified PvTRAg proteins were multimerised on Nile Red (NR) coated streptavidin beads (**Figure 2A**). Reticulocytes enriched using anti-CD71 magnetic beads (**Supplementary Figure 5**) were mixed 1:1 with mature erythrocytes and then incubated with the PvTRAg-NR beads. After incubation and washing, the mixture was stained with Thiazole Orange (TO), which binds the RNA present in reticulocytes but not mature erythrocytes and allows detection of binding to reticulocytes and/or erythrocytes by flow cytometry, double gating with NR and TO. Pfs25, a sexual stage protein from *P. falciparum* [25-27] with no known red cell binding activity was used as a negative control, while the reticulocyte-specific CD71-interacting domain of PvRBP2b_169-813_[8] acted as a positive control. Of the 15 PvTRAgs tested, five showed binding significantly above the levels of the Pfs25 negative control, with binding by PVP01_0000100 (>60%) and PVP01_0503700 (>20%) being the most significant with clear preference for reticulocytes over erythrocytes (**Figure 2B**), similar to the preference of PvRBP2b. (**Figure 2B**; gating strategy shown in the **Supplementary Figure 6**).

**Figure 2:**
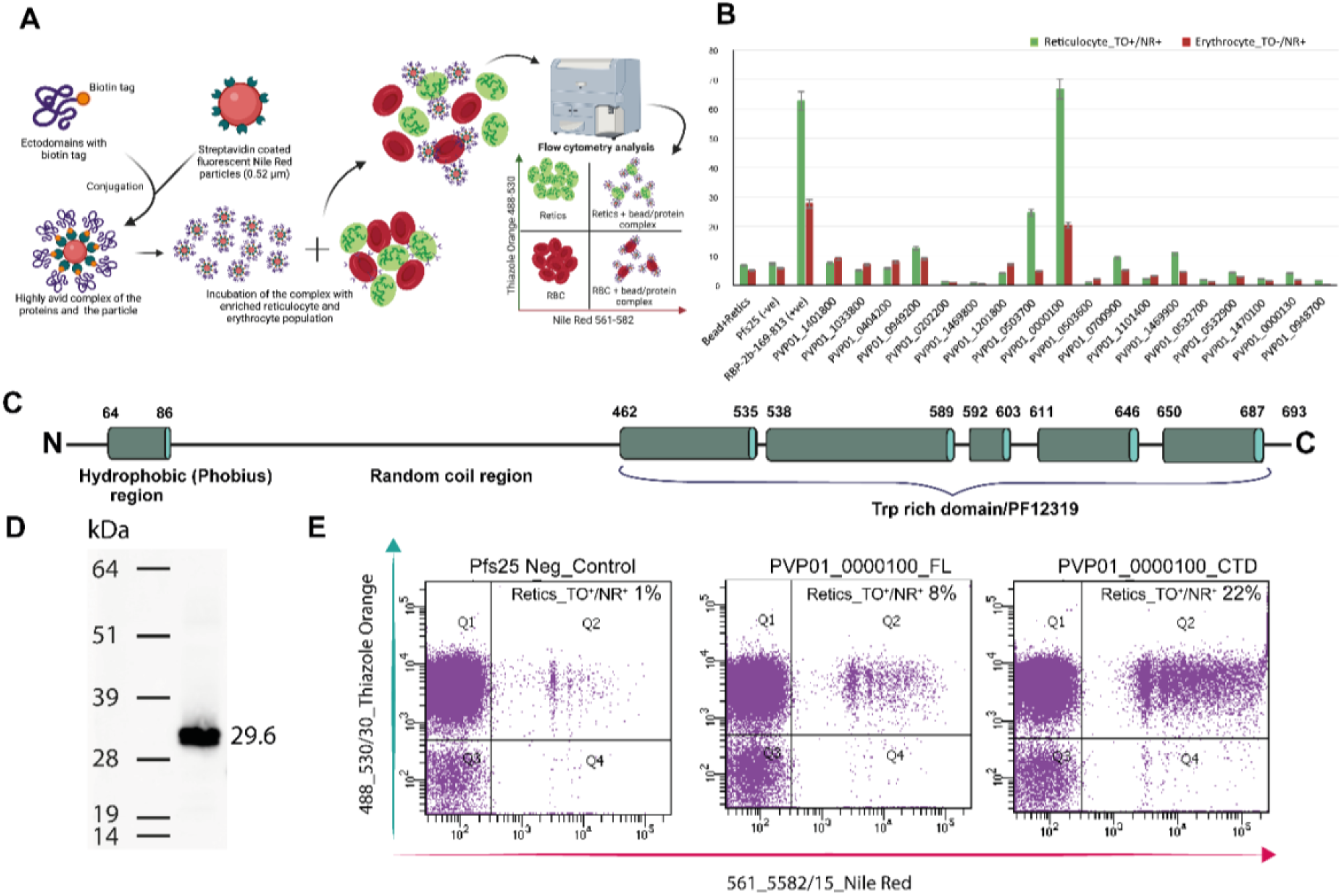
Screening PvTRAgs for erythrocyte and reticulocyte binding activity. **A) Scheme of red blood cell binding assay**. Affinity purified biotinylated recombinant proteins were multimerised on streptavidin coated beads and incubated with a mixture of erythrocytes and reticulocytes. Reticulocytes were labelled with Thiazole Orange (staining mRNA), before binding was assessed using flow cytometry, with TO excited with a 488nm laser and Nile Red with a 561nm laser. Erythrocytes are devoid of any genetic material, so are not labelled with TO and in the lower half of the plots. Any binding with protein-labelled beads will cause a shift in the population of cells towards upper right (reticulocyte, green) or lower right (erythrocyte, red). **B) Binding of PvTRAgs to erythrocytes and reticulocytes**. Bar graph showing the binding of the multimerised PvTRAgs to reticulocytes and erythrocytes. PvRBP2b_169-813_ previously reported to have binding activity against reticulocytes was used as positive control, while Pfs25, a sexual stage antigen, was used as a negative control for this assay. Data are graphed as mean ± S.D. n=3. **C) PVP01_0000100 domain prediction**. Schematic diagram of PVP01_0000100 secondary structure prediction using NetSurfP 3.0 and Phobius identies a long N-terminal random coil region and a C terminal trp-rich region which is predicted to possess an extensive helical secondary structure. The alpha helices are depicted as deep green coloured barrel while the random coil regions as black strand. **D**) **Purification of PVP01_0000100 C-terminal domain (CTD)**. SDS PAGE showing purified PVP01_0000100 CTD, encompassing amino acid D459-693L, as single protein band with the correct mass (29.6 kDa) following two-step (affinity and size exclusion) purification. **E) The PvP01_0000100 CTD contains reticulocyte binding properties**. The reticulocyte binding activity of the PVP01_0000100 CTD and its full-length counterpart was assessed in flow cytometry as above.

PVP01_0000100 and PVP01_0503700 are close to each other in the phylogenetic tree (**Figure 1A**) and share 34% amino acid identity across the full-length proteins, and 48% identity in their C-terminal Tryptophan/Threonine-rich domains. Secondary structure prediction for PVP01_0000100 using NetSurfP 3.0 suggests that the Tryptophan/Threonine-rich domain possesses a high α-helical content (**Figure 2C**). To test which domain was responsible for reticulocyte binding, the C-terminal region was expressed alone (**Figure 2D**) and assessed along with its full-length counterpart. The C-terminal Tryptophan/Threonine-rich domain showed an even higher level of binding towards reticulocytes (**Figure 2E**), establishing that the reticulocyte binding activity is mediated by this domain.

### The PVP01_0000100 C-terminal Tryptophan/Threonine-rich domain adopts a Bin/Amphiphysin/Rvs (BAR) domain fold

To establish how the C-terminal Tryptophan/Threonine-rich domain of PVP01_0000100 mediates reticulocyte binding we determined the X-ray crystal structure of this domain. The structure was refined to 1.45 Å resolution and had one molecule in the asymmetric unit (**Figure 3A**). Consistent with the NetSurfP predictions, the structure is composed primarily of α-helices that are arranged into an extended three-helical bundle with small helical domains at each end. The conserved tryptophan residues that define the domain are distributed along the length of the helical bundle and are all buried within the fold, contributing to the arrangement of the long helices. Since experimental determination of this structure, new methods for prediction of 3D protein structures have been accelerated via use of deep-learning strategies as exemplified by AlphaFold2 (AF2)[28]. AF2 predictions of the PVP01_0000100 Tryptophan/Threonine-rich domain structure closely resemble the structure we experimentally determined, with RMSD 1.56 Å over 213 C-alpha atoms (**Supplementary Figure 7A)**. The accuracy of the predicted AF2 structure therefore allows for comparison of our experimental structure with AF2 predictions for the Tryptophan/Threonine-rich domain of TRAgs from other *Plasmodium* species. The quality of the predicted models was assessed by analysis of the pLDDT plots which showed high confidence scores for the helical regions of these predicted models (showed inset), (**Supplementary Figure 7C**). Superposition of PVP01_0000100 Tryptophan/Threonine-rich domain with its closest orthologue in *P. knowlesi* (PKNH_1300500, RMSD 1.83 Å over 211 C-alpha atoms, seq identity 77.4%) and a much more distant orthologue in *P. falciparum* (PF3D7_0102700, RMSD 2.87 Å over 179 C-alpha atoms, seq identity 30.2%) suggests strongly that the three-helical fold we experimentally determined is highly conserved across *Plasmodium* species (**Supplementary Figure 7B**).

**Figure 3:**
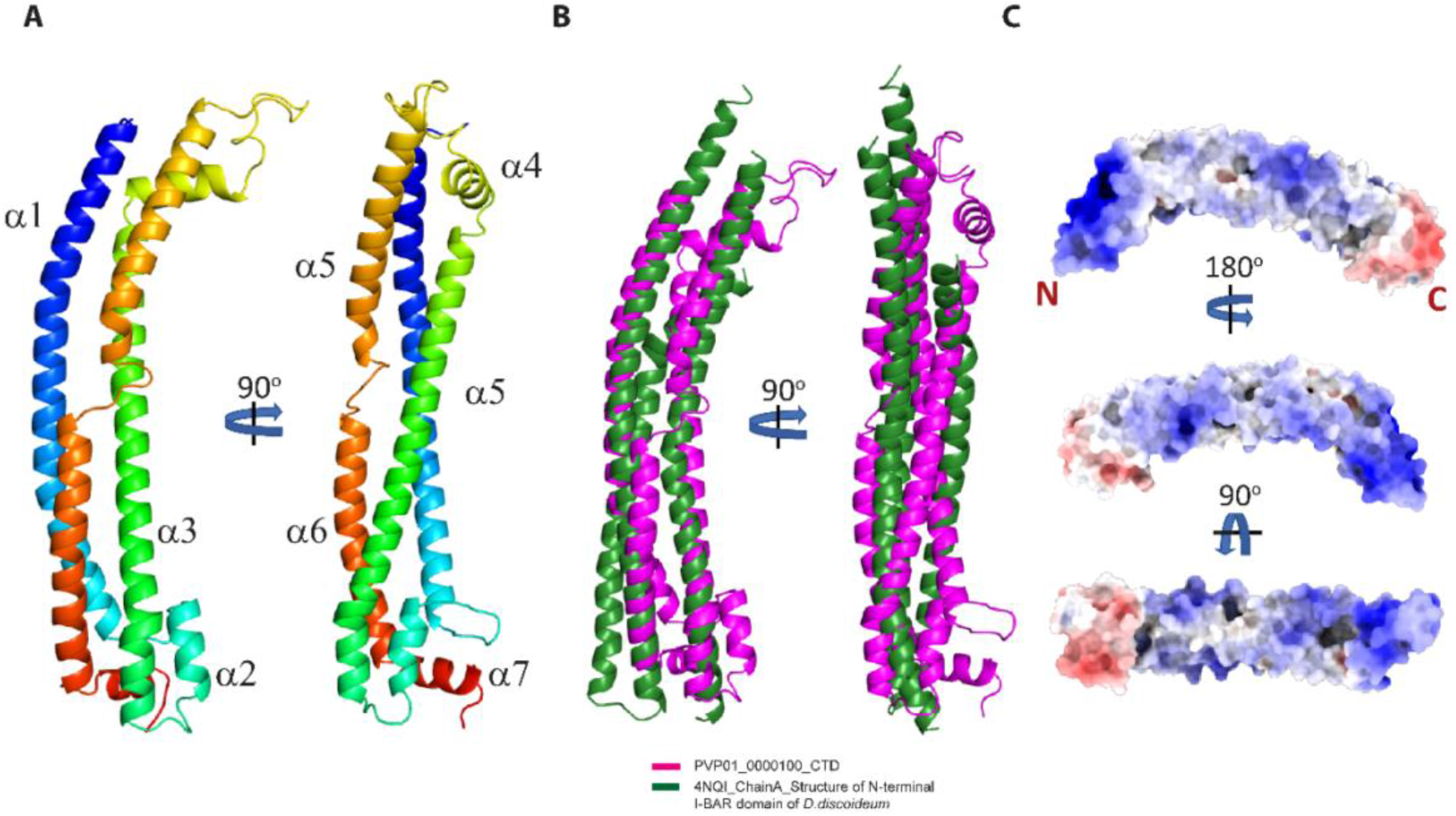
Structure of the PVP01_0000100 CTD and its homology with other BAR domain protein. **A) Structure of the CTD of PVP01_0000100**. Ribbon diagram of the CTD of PVP01_0000100 coloured from blue (N-terminus) to red (C-terminus). Two orientations are displayed (rotated by 90°) demonstrating the extended, slightly bent, three-helical bundle structure **B) Structural homology of the C terminal TRAg domain to a BAR domain protein of *Dictyostelium discoideum***. Superposition of the CTD of PVP01_0000100 (magenta) with the BAR domain structural homologue from *Dictyostelium discoideum* (PDB ID 4NQI, green) showing the similarity between the fold of these two domains. **C) Electrostatic charge distribution on the TRAg domain surface**. Electrostatic potential analysis of the surface of PVP01_0000100 identifies positively charged patches at the N-terminal end and along the surface including the concave side of the protein. The small domain at the C-terminal end of the TRAg domain possesses a negatively charged surface.

The structure of the Tryptophan/Threonine-rich domain of PVP01_0000100 was compared to other known protein folds using DALI[29]. This identified similarity to several membrane-binding proteins containing Bin/Amphiphysin/Rvs BAR domains (PDB ID 4NQI with a DALI Z-score of 7.2) (**Figure 3B**). BAR domains possess extended helical bundles with a bent shape that can bind lipids and in some cases reshape membranes. Typically, BAR domains interact with negatively charged lipids on the plasma membrane via positively charged patches on the protein surface[30-33]. Analysis of the surface charge of the PVP01_0000100 revealed extended positively charged patches mainly on the N-terminal and concave side of the protein surface (**Figure 3C)**, similar to the crescent shape and surface charge of BAR domains, and suggesting suggests a potential functional role in binding lipids.

### The PVP01_0000100 Tryptophan/Threonine-rich domain binds to sulfatide

In order to carry out comparative *in vitro* lipid-binding assays, full-length PVP01_0000100 (FL) protein, PVP01_0000100 CTD and a negative control protein Pfs25 were all purified (**Figure 4A**). As PVP01_0000100 is expressed on the merozoite surface, the lipid species it is likely to interact with will be those present in the outer leaflet of the host plasma membrane. We therefore performed lipid-binding assays with binding strips containing a range of sphingolipids, which are abundant in the outer leaflet of cell surface membranes. PVP01_0000100 CTD showed specific and clear binding to sulfatide (**Figure 4B**), an anionic glycosphingolipid found on the outer leaflet of multiple cell types, including red blood cells[34, 35]. The interaction with sulfatide was confirmed using giant multilamellar liposomes composed of 48% PC, 2% Rhod-PE and 50% sulfatide. These liposomes were incubated with purified protein, pelleted via centrifugation, washed, and run on an SDS PAGE. Both PVP01_0000100 FL and CTD bound more strongly to liposomes containing 50% sulfatide than they did to liposomes composed of only 98%PC and 2%Rhod-PE, whereas Pfs25 didn’t show any significant binding to either (**Figure 4C**).

**Figure 4:**
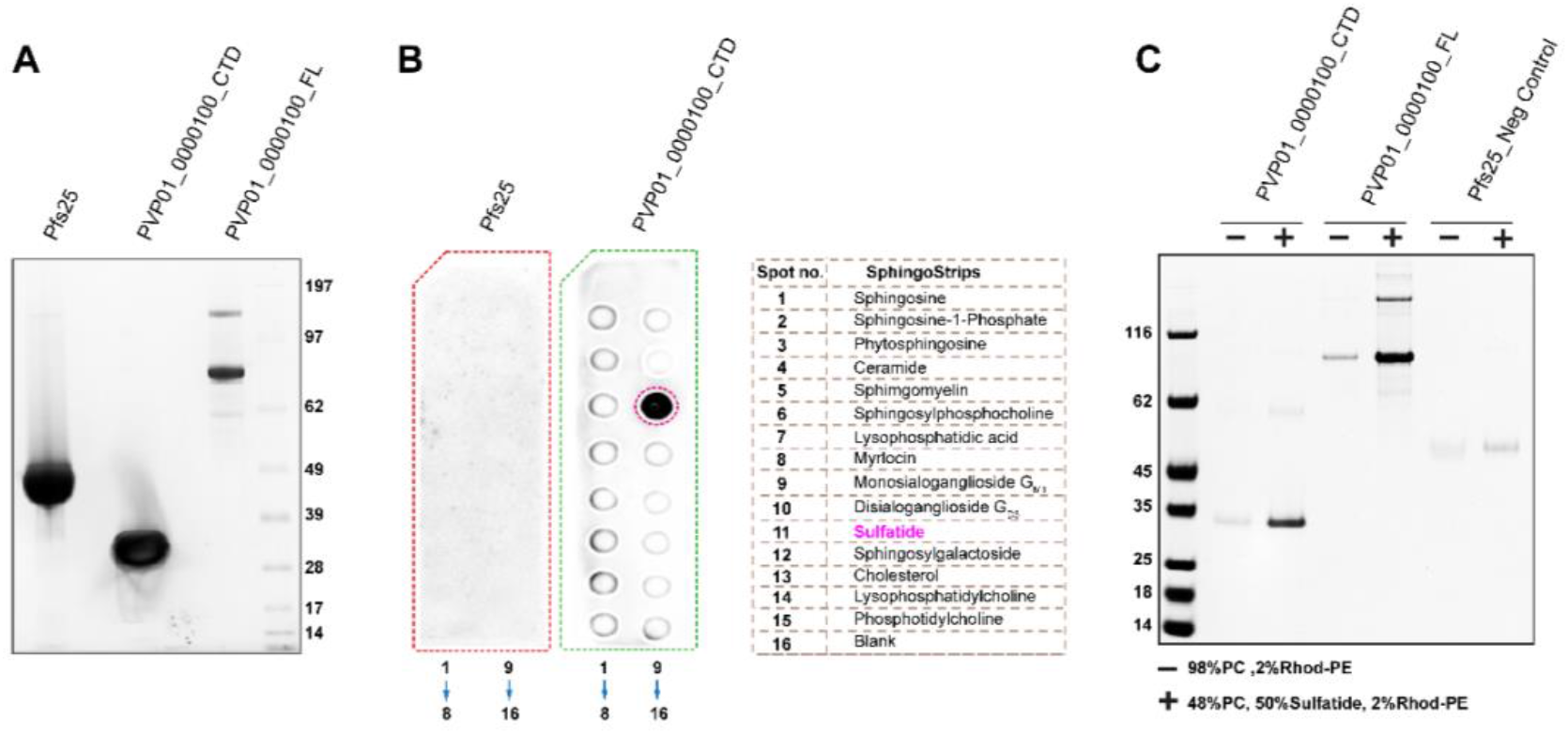
Lipid binding properties of PVP01_0000100. **A) Purity of expressed recombinant proteins**. SDS-PAGE of purified PVP01_0000100 full length (FL), PVP01_0000100 C-terminal domain (CTD) and Pfs25 following affinity and size exclusion chromatography demonstrating the high purity of all three proteins. **B) Binding to sphingolipids**. PVP01_0000100 CTD and Pfs25 were incubated with sphingostrips and binding detected using HRP conjugated anti-His antibody (1:500) and chemiluminescent substrate. 1µg (0.03µM) of proteins was used in this assay, with Pfs25 acting a negative control. The spot numbers and orientation are shown under the blot, and indicate PVP01_0000100 CTD binding strongly and specifically to sulfatide (spot no.11, magenta circle). **C, Protein-liposome binding assay**. Liposomes composed of PC only (-) or 48:50:2 PC:sulfatide:Rhod-PE (+) were assessed for binding of PVP01_0000100 FL, CTD and Pfs25 at 1µM. The band intensity is increased in the presence of sulfatide for both CTD and FL PvP01_0000100 proteins, while the negative control, Pfs25 did not significantly bind liposomes.

These two independent assays confirm the lipid-binding properties of the PvP01_0000100 Tryptophan/Threonine-rich domain that were suggested by structural homology to BAR domains. This is the first confirmed molecular function for a *Plasmodium* Tryptophan/Threonine-rich domain and given the likely conservation of this fold (**Figure 3B**), may be a universal feature of all *Plasmodium* TRAgs. Interestingly, comparison of the surface charge of TRAgs from different *Plasmodium* species revealed variation in the surface charge distribution, suggesting that each member of the TRAg family may have specificity towards different lipids (**Supplementary Figure 8**).

### The *P. knowlesi* homologue of PVP01_0000100 plays a role in reticulocyte invasion

*P. vivax* parasites cannot be cultured *in vitro*, which means the function of PVP01_0000100 cannot be directly tested in this species. To test the implications of sulfatide binding we therefore generated a knockout line of the PVP01_0000100 orthologue (PKNH_1300500) in *P. knowlesi*, a close phylogenetic relative of *P. vivax* which can be cultured in human erythrocytes [36]. Purified PKNH_1300500 bound both sulfatide and reticulocytes, just like PVP01_0000100 (**Supplementary Figure 9**). Deletion of PKNH_1300500 was carried out using a CRISPR-Cas9 two-plasmid approach, resulting in deletion of the entire gene and replacement with a GFP expression cassette (**Figure 5A**; see Methods for details). After transfection and selection, genotyping and GFP fluorescence both revealed a mixed population of wildtype and edited parasites, from which knockout lines were isolated by dilution cloning **(Figure 5B)**. We have previously shown that anti-PVP01_0000100 antiserum detects merozoite surface staining in wild type *P. knowlesi* schizonts (**Supplementary Figure 2)**. The fluorescence intensity of this staining was reduced by >80% in the PKNH_1300500 knockout line (**Figure 5C**), confirming gene deletion. The weak residual staining suggests that there might be some low-level of cross-reaction of anti-PvP01_0000100 antisera with PkTRAgs other than its closest homologue PKNH_1300500, although previous tests showed only limited cross-reaction (**Supplementary Figure 1**).

**Figure 5:**
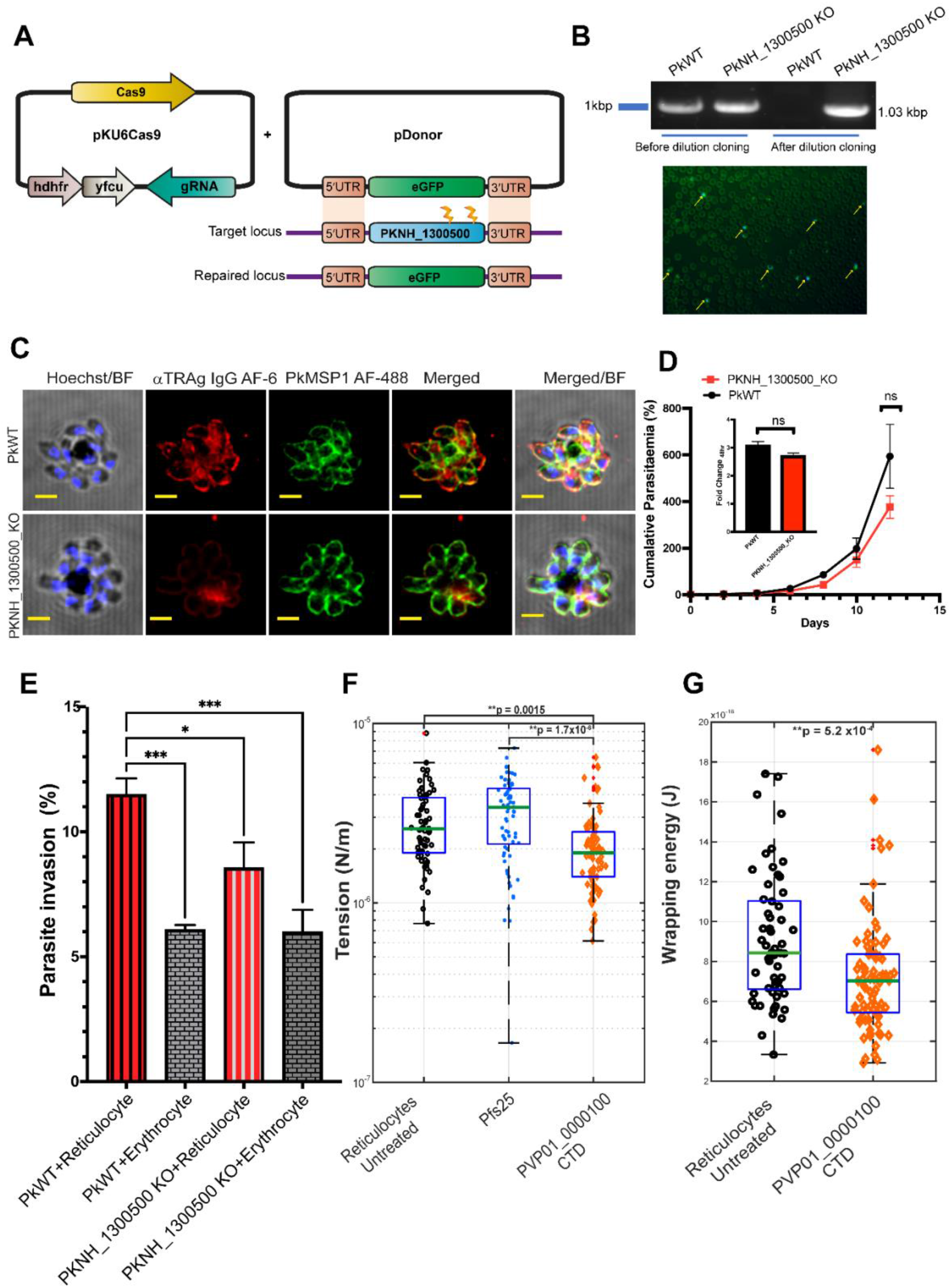
Gene deletion in *P. knowlesi* reveals a function for PKNH_1300500 in reticulocyte invasion. **A) CRISPR-Cas9 editing strategy**. The homologous repair template used for gene editing of the PVP01_0000100 orthologue in *P. knowlesi* (PKNH_1300500) contained an eGFP expression cassette flanked by 5’ and 3’ UTR homology regions of 800 bp each. This construct was co-transfected with a plasmid expressing both Cas9 and two guide RNAs positioned near the 3’ end of the gene. **B) Confirmation of knockout by PCR and GFP fluorescence**. After transfection and drug (100 nM pyrimethamine) selection, genotyping PCR revealed a mixed population of PkWT and PKNH_1300500 parasites, as indicated by PCR specific for the wildtype (WT) and knockout (KO) locus. A pure population of the edited line was obtained by dilution cloning, with no WT genotyping visible (lane 3). Successful editing was also assessed using GFP expression (yellow arrows showing the positively edited cells) under fluorescent microscope. **C) Immunolocalisation using anti-PVP01_0000100 antisera in WT and KO *P. knowlesi* parasites**. Rabbit polyclonal IgGs raised against PVP01_0000100 was used to detect its orthologue PKNH_1300500 in *P. knowlesi*. Alexa Fluor 488 goat anti rat secondary antibody was used in combination with Alexa Fluor 633 goat-anti rabbit secondary antibody. Top panel is the *P. knowlesi* wild type population and bottom panel is PKNH_1300500 gene edited population. PkMSP_1-19_ antibody (raised in rat) was used as merozoite surface marker and Hoechst was used to stain the parasite nucleus. **D) WT vs KO growth rate comparison**. Growth rate of the PkWT and PKNH_1300500 KO line was assessed for 14 days, with parasites split every 2 days and reset to 1% parasitemia. Cumulative parasitaemia was calculated by multiplying parasitemia values detected with flow cytometry with the cumulative split values. Data points represent two independent biological replicate and graphed as mean ± S.D. p>0.05 (unpaired two-sided t-test) **E**) **Invasion quantification**. Invasion was quantitated using flow cytometry, with parasites detected using SBYR green staining, and erythrocytes and reticulocytes distinguished using anti-CD71. The percentage of reticulocytes or erythrocytes that were invaded is shown (mean ± S.E.M) and the data combines three biological replicates, one containing technical duplicate wells, and the other two technical triplicate wells. P*<0.05, P**<0.01, P***<0.001 (one-way Anova test) **F) Effect of PVP01_0000100 on reticulocyte biophysical properties**. Membrane tension was measured using flickering spectroscopy for CD71+ reticulocytes in the absence of additional protein (untreated), and or incubated with 100 µg/ml proteins (Pfs25 and PVP01_0000100). Number of cells per sample: untreated n = 65, Pfs25 n = 73, and PVP01_0000100 n = 79. Pairwise comparisons between groups used Tukey’s honest significant difference (HSD) test and Bonferroni correction. **G) Analysis of the wrapping energy of reticulocytes**. Wrapping energy was calculated using MATLAB. Data points for untreated are 57 and PVP01_0000100 sample are 77. Pairwise comparisons between groups used Tukey’s honest significant difference (HSD) test and Bonferroni correction.

To assess the impact of PKNH_1300500 deletion, growth assays were performed in duplicate for 14 days (c. 14 cycles, as *P. knowlesi* has a 27-hour cycle). A slight reduction in cumulative growth of the PKNH_1300500 KO line compared to wildtype *P. knowlesi* was observed, but this was not statistically significant (**Figure 5D**). As both PVP01_0000100 and PKNH_1300500 bind preferentially to reticulocytes, we carried out invasion assays comparing the ability of the KO line to invade both reticulocytes and mature erythrocytes. Late stage parasites were incubated with a mixture of erythrocytes and reticulocytes and invasion was quantitated using flow cytometry, with anti-CD71 staining distinguishing between erythrocytes and reticulocytes, and invasion events quantitated using SYBR Green to label parasite DNA **(Supplementary Figure 10)**. Using a CD71 positive cutoff to define reticulocytes has a risk of false negatives, as some older reticulocytes may have very low CD71 levels [37], but it reduces the risk of false positives from low levels of contaminating erythrocytes that would also be CD71 negative. Wildtype *P. knowlesi*, while not restricted to reticulocytes like *P. vivax*, does have a clear preference for these immature red cells [38]. The preference was observed as expected in wildtype *P. knowlesi*, whereas in PKNH_1300500 knockout parasites the preference was significantly reduced (**Figure 5E**). The impact of PKNH_1300500 deletion specifically on reticulocyte invasion likely explains the lack of difference in cumulative growth rate between KO and wildtype lines, as under standard growth conditions, parasites are cultured only in the presence of mature erythrocytes, with few if any reticulocytes present (as these mature rapidly in culture).

To explore the mechanism by which PVP01_0000100 might play a role in invasion, we incubated purified PVP01_0000100 CTD with reticulocytes and assessed the impact on reticulocyte biophysical properties using flickering spectroscopy[39]. The PVP01_0000100 CTD led to a significant decrease in membrane tension, which in *P. falciparum* has previously been shown to be directly linked with invasion efficiency, with lower tension erythrocytes being more accessible for invasion than higher tension erythrocytes [40]. Purified Pfs25 used at the same concentration had no effect (**Figure 5F**). Tension is thought to affect invasion through its impact on the membrane wrapping energy, essentially the amount of energy required for the parasite to wrap the cell membrane around itself during the invasion process [41]. Addition of PVP01_0000100 reduced the wrapping energy compared to untreated reticulocytes (**Figure 5G**), suggesting a model by which PVP01_0000100 binding to sulfatide modulates invasion by decreasing the energy required for merozoites to enter their host.

## Discussion

Tryptophan Rich Antigens (TRAgs) have been studied in multiple *Plasmodium* species and have been suggested as potential vaccine targets, but their molecular function has never been established, and the previous literature around them is sometimes contradictory, suggesting a new systematic analysis is required. A comprehensive search and annotation of TRAgs from *P. vivax, P. cynomolgi* and *P. knowlesi* revealed that half of the genes are predicted to contain signal peptides, but the other half are not. From a cell biological and evolutionary perspective, it would be unusual for the same protein family to contain members both with and without signal peptides, as secreted and cytosolic proteins generally have quite different functions. A more prosaic, technical explanation therefore seems likely. The genomes of *Plasmodium* species are all biased towards a high AT content and have patterns of codon usage that are unusual compared to other eukaryotes [42, 43]. This can affect the efficiency of genome annotation for protein features such as signal peptides, which use domain prediction programmes that are usually trained on genomes from humans or model eukaryotes. We observed several TRAgs had SPs annotated by some prediction programmes but not others, and all contained a hydrophobic sequence at their extreme N-terminus, regardless of whether it is formally annotated as an SP. Immunofluorescence assays using antibodies raised against four *P. vivax* TRAgs, two with predicted SPs and two without, revealed that all four co-localised with a merozoite surface marker, so all must enter the secretory pathway. Based on this we believe that it is likely that the genome annotation of this gene family is incomplete and that all TRAgs are likely secreted, regardless of whether the hydrophobic region at their N-terminus is formally annotated as a signal peptide. This emphasises the importance of validating annotation and exploring the use of multiple annotation servers when analysing *Plasmodium* genes, particularly those of unknown function.

The presence of multiple PvTRAgs on the merozoite surface, confirmed by this and previous studies [24], suggests a role for some family members in erythrocyte and reticulocyte binding. Using a new bead binding assay we observed strong binding for PVP01_0000100 and PVP01_0503700 with a preference for reticulocytes in both cases, which has not been reported previously in studies of other PvTRAgs [12, 13]. PKNH_1300500, the *P. knowlesi* homologue of PVP01_0000100, also showed reticulocyte binding activity, although this was weaker than PVP01_0000100, perhaps due to the weaker expression of this construct which may affect the conjugation efficiency with the beads. Solving the crystal structure of the Tryptophan/Threonine-rich domain of PVP01_0000100 suggested a hypothesis for the molecular mechanism of this binding activity. The three helical bundle structure of this domain has homology with the BAR domain superfamily, which is characterized by α-helical coiled coils that form a crescent shape with positively charged patches on the concave side, which interacts with phospholipid membranes[30, 44]. BAR domains are often involved in membrane curvature, for example by interacting with negatively charged membranes and contributing to membrane bending during endocytosis[45], and can also serve as membrane-binding scaffolds for other proteins, thereby participating in signalling pathways[46]. The PVP01_0000100 Tryptophan/Threonine-rich domain has positively charged patches on the concave side (driven by high lysine content, 18%), and lipid dot blot and liposome binding assays confirmed that both it and its orthologue in *P. knowlesi* (PKNH_1300500) binds specifically to the anionic lipid sulfatide. Based on what is known about BAR domain lipid binding, there is unlikely to be a specific sulfatide binding pocket within the PVP01_0000100 Tryptophan/Threonine-rich domain, with binding instead mediated by charge interactions across the curved surface.

Sulfatides (also known as 3-O-sulfogalactosylceramides, sulfated galactocerebrosides, or SM4) are sphingolipids that participate in a wide range of cellular processes including protein trafficking, cell adhesion and aggregation, axon-myelin interactions, neural plasticity, and immune responses, among others. They are particularly enriched in the myelin sheath surrounding nerve cells but are also found in the extracellular leaflet of the plasma membrane of most eukaryotic cells including red blood cells. Sulfatides have previously been reported to play a role in malaria infection, specifically during the invasion of hepatocytes where circumsporozoite protein (CSP) has been reported to bind to sulfatide [47] [48] Sulfatide binding has not previously been implicated in erythrocyte invasion in any *Plasmodium* species, but deletion of PKNH_1300500, which binds sulfatide just like its *P. vivax* orthologue, affects both parasite growth and invasion, suggesting a direct link between sulfatide binding and red blood cell invasion for the first time. The homology between the PVP01_0000100 Tryptophan/Threonine-rich domain fold and the BAR domain suggests a clear hypothesis for how TRAgs might promote invasion. BAR domain binding to lipid moieties can cause local curvature in membranes, driving processes such as endocytosis [45]. TRAg binding to lipids on the surface of erythrocytes could therefore cause local membrane remodelling, potentially affecting biophysical properties and aiding the invasion process, which requires significant reshaping of the erythrocyte membrane in order for the merozoite to enter. In keeping with this hypothesis, addition of purified PVP01_0000100 to reticulocytes reduced their surface tension **(Figure 5G)**. Lower surface tension has recently been associated with increased invasion efficiency [40], supporting the hypothesis that TRAg-lipid interactions could directly promote invasion in reticulocytes. This interaction is clearly not absolutely essential for invasion, as PKNH_1300500 knockout parasites can still invade both erythrocytes and reticulocytes, though at a reduced rate in reticulocytes. This is likely due to the large number of genes within the TRAg family, with other PkTRAgs, presumably binding to either sulfatide or other lipids, providing partially overlapping functions. Whether the sulfatide binding preference of PVP01_0000100 also explains its reticulocyte binding preference is not clear, as we were unable to find any data quantifying the abundance of sulfatide on reticulocytes compared to erythrocytes. It is worth noting though that CD71^+^ reticulocytes have a c. 30% larger surface area compared to mature erythrocytes, and membrane lipids vesiculate out during the process of RBC maturation, suggesting that many lipids, including sulfatide, are likely to be more abundant in reticulocytes [37, 49].

We have used the localisation and structure of a specific merozoite surface localised PvTRAg, PVP01_0000100, to generate and test a specific hypothesis – that this protein binds lipids and thereby promotes the invasion of erythrocytes by *P. vivax* merozoites. The obvious question is whether this hypothesis holds true across all TRAgs within *P. vivax*, or indeed for all TRAgs across the *Plasmodium* genus. AlphaFold2 predicts that the structure we experimentally determined for the PVP01_0000100 Tryptophan/Threonine-rich domain is likely to be conserved across *Plasmodium* TRAgs, despite sequence conservation levels <50%. This strongly suggests that all will possess a BAR domain-like fold, and hence all may bind lipids. However, the low sequence homology and different surface charge patterns seen between different *Plasmodium* Tryptophan/Threonine-rich domain models suggests that it is very possible that individual TRAgs bind to different lipids. It is also worth noting that other *P. vivax* TRAgs have been suggested to be localised not on the merozoite surface but are exported into the infected erythrocyte cytosol, where they associate with Maurer’s Clefts or other membrane structures [50]. Similarly, a *P. berghei* TRAg has recently been shown to be found inside infected erythrocytes and proposed to be involved in membrane remodelling [51] These observations are entirely in keeping with our hypothesis that all TRAgs have lipid binding properties – they simply may do so with different lipid specificities, and in different intracellular locations. This work has established the first molecular function for the TRAg family and provides clear hypotheses to test in the further exploration of this unique *Plasmodium* protein family.

## Material & Methods

### Phylogenetic tree generation

The amino acid sequences of TRAgs from *P. vivax* (38), *P. cynomolgi* (33), *P. knowlesi* (29) were obtained from PlasmoDB [20, 52]. The full length amino acid sequences of these TRAgs were aligned using Clustal Omega [53] and trees generated using PhyML [54]. Phylogenetic trees were constructed using maximum likelihood method [55]. The evolutionary distances were computed using Jones-Taylor-Thornton Matrix model with a bootstrap test of 100 replicates [56] and remove gap from alignment option.

### Gene synthesis and Protein expression

The region corresponding to the entire extracellular domain of 33 *P. vivax* Tryptophan rich antigens were determined using signal peptide [57] and transmembrane [58] prediction servers. Five of the 38 total *P. vivax* TRAGs were omitted because they were >200 kDa in size; proteins above this size are usually not expressed well in HEK239E cells[23]. To prevent the aberrant glycosylation of the proteins in mammalian cells all potential N-linked glycosylation sites (N-X-S/T/P), which are non-functional in *Plasmodium* parasites, were mutated by substituting alanine for serine/threonine/proline. The designed constructs were synthesised (TWIST biosciences, UK) with codons optimised based on the human genome. The protein coding region was flanked by *Not*I and *Asc*I site and subcloned in a pTT3 expression plasmid between a 5’ mouse variable κ light chain signal peptide [59] [23], an enzymatic biotinylation sequence and a hexa-His tag. The synthesised plasmids were amplified, purified, and diluted to 1 mg/ml then mixed in a 1:2.5 ratio with Polyethylenimine (PEI 40 kDa) used as a transfectant (for transfection in 50 ml, 25 μl plasmid (1 mg/ml) in 2.5 ml and 62.5 μl PEI in 2.5 ml freestyle media were mixed). To biotinylate the recombinant proteins, 1.25 μl *E. coli* BirA ligase plasmid was co-transfected with the plasmid of interest. The DNA/PEI mixture was then incubated for 8min and added to HEK 293E cells. The cells were cultured in Freestyle 293 media (Invitrogen, USA) supplemented with Geneticin (Gibco, USA) and 1% foetal bovine serum (Gibco, USA) and were maintained in a shaker flask at 125 rpm, 37°C, 5% CO_2_ and 70% relative humidity.

### Western Blotting

In the HEK239E cell/pTT3 system recombinant proteins are secreted into the culture supernatant. To confirm the expression of the recombinant proteins, 10 μl of culture supernatant was collected from each transfection and mixed with 3.5 μl of NuPAGE 4X LDS sample buffer and 1.5 μl of sample reducing agent (Invitrogen, USA). The samples were heated in 80°C for 10 mins and resolved using SDS PAGE (4-12% Polyacrylamide gel) followed by Western blotting to Nitrocellulose membrane (Amersham), with transfer carried out at 100 V for 60 mins at 4°C. Membranes were blocked in 3% BSA in TBST (Tris buffer saline with 0.05% Tween20) for 2 hrs at room temperature and probed using HRP conjugated anti-His antibody (Proteintech, Ptglab, USA, 1:20,000) overnight at 4°C. After 3 × 10 min washes with TBST buffer, expressed proteins were detected using ECL chemiluminescent substrate (GE Healthcare, USA) in Biorad ChemiDoc MP™ Imaging System. The volume of culture required was optimised for each construct based on the expression level in the blot.

To assess cross reactivity of PVP01_0000100 with other TRAgs, an equal amount (5 μg) of 14 purified PvTRAgs were run on a single gel, and probed with polyclonal antibodies raised against PVP01_0000100. Pfs25 was used as control, as it shares the biolinker hexa-His tag that is common to all the constructs. The protocol for the blotting was largely as above, except blocking was carried out in 5% milk TBST at room temperature (RT) for 2 h followed by overnight incubation at 4°C with Rabbit IgGs against the respective TRAgs (1:20,000). Blots were washed for 3 × 10 mins with TBST, then incubated with HRP conjugated anti rabbit secondary antibody (1:20,000) for 45 mins. Following three further washes, antibodies were detected using ECL chemiluminescent substrate (GE Healthcare, USA) in a Biorad ChemiDoc MP™ Imaging System.

### Protein purification

To purify proteins from supernatant, HEK 293E cells were removed by pelleting at 3000g for 20 mins followed by filtering (0.22 µm, Corning). Culture supernatant was adjusted to 25 mM Imidazole and 300 mM NaCl, then loaded on to a 1 ml HiTrap Ni column (Cytiva) preequilibrated with binding buffer (40 mM Imidazole, 20 mM Sodium Phosphate, 500 mM NaCl, pH 7.4) at 1 ml/min flow rate. After completion of sample loading, the column was washed with 5 column volume (CV) of binding buffer to release any non-specifically bound protein. Elution was performed using step gradients with elution buffer containing 400 mM Imidazole, 20 mM Sodium Phosphate, 500 mM NaCl, pH 7.4. The eluted fractions containing recombinant protein were collected and pooled together for buffer exchange in PBS and concentration in 10kDa Vivaspin Ultracel concentrators. The final concentrations were measured by Nanodrop OneC UV-Vis Spectrophotometer (Thermo Fisher) and kept in 4°C until further use.

### Immunisation and IgG purification

Polyclonal IgGs were developed in New Zealand White Rabbits (Eurogentec, Belgium) as per the company’s protocol. In brief, 0.8 mg of purified recombinantly *P. vivax* TRAg protein was injected in two rabbits (0.4 mg each) with 100ug/immunisation/rabbit. Freund’s complete adjuvant was used for first immunisation followed by Freund’s incomplete adjuvant in booster doses given on days 14, 28 and 58. The final bleed was collected on day 87.

Total IgG was purified from the final bleed rabbit sera using Protein G Gravitrap (GE Healthcare, Cytiva). The sera were diluted as 1:5 in binding buffer (20 mM Sod. Phosphate, pH 7.4). The purification was done using Ab purification kit and the protocol was followed as per the kit’s manual. In brief, the diluted sera were loaded in the preequilibrated Gravitrap column and the eluant reloaded three times to enrich the binding and the beads washed with 10 ml of binding buffer. The IgG was finally eluted with 5 ml of elution buffer (0.1 M glycine-HCl, pH 2.7) poured into the column and collected directly in a 10 kDa centrifugal filter unit (Millipore) containing 0.225 ml of neutralisation buffer (1 M Tris-HCl, pH 9.0). The IgGs were buffer exchanged three times in RPMI 1640 media (Gibco) for any follow up assay. The final concentration of the purified IgG was determined using a Nanodrop one UV-Vis Spectrophotometer (Thermo Fisher).

### Reticulocyte Isolation

#### Enrichment of CD71+ reticulocytes

Fresh peripheral whole blood (withdrawn within 48 hours) provided by NHS Blood and Transplant (ethical approval University of Cambridge REC HBREC.2019.40 and NHS REC 20/EE/0100). Whole blood was centrifuged for 7 mins at 1500 g and the plasma and the white cell layer aspirated, then resuspended with equal volume of phosphate-buffered saline buffer, PBS (Gibco, UK). The solution was passed through a Plasmodipur filter to remove leukocytes (EuroProxima, The Netherlands) and washed twice with PBS at 1000 g for 10 mins. The leukocyte-depleted red blood cell pellet was then resuspended in 10 ml PBS and 5 ml were carefully layered on 6 ml of 70% (v/v) isotonic Percoll cushion (GE Healthcare, UK). After centrifugation at 1200 x g for 15 mins, brakes off, a thin band(s) of reticulocytes was collected at the Percoll interface and spun down at 300 g for 10 mins. All the following steps were carried out at 4°C to preserve reticulocytes. Microscopic counting of reticulocytes stained with new methylene blue (NMB, Sigma-Aldrich, UK), and the Percoll separation achieved purity of 7% - 20% reticulocytemia, depending on the blood sample, as previously reported [60]. To purify immature CD71^+^ at purities up to 90%, we used CD71 magnetic MicroBeads (Miltenyi Biotec, UK) and followed the company protocol. We confirmed the high purity through sub vital staining (NMB) and microscopy. Reticulocytes were stored at 4°C and used for binding experiments within 24 hs.

#### Reticulocyte binding assay

Purified biotinylated proteins were multimerised on streptavidin coated Nile Red fluorescent beads (Spherotech Inc, 0.52 μm, 0.1% w/v, binding capacity 1.65 nM Biotin-FITC to 1 mg beads). The beads were sonicated to disperse them evenly using a bath sonicator for 10 mins with a pulse duration 30 secs and 10 secs interval between each pulse, all carried out at 4°C. The sonicated beads were then blocked using 0.2% BSA buffer (in PBS). In a 96 U bottom plate (Nunc 165306, Thermo), 1 μl bead was incubated with 1 μg protein per well for 45 min at 4°C. Unbound protein was then removed by washing the beads by centrifugation at 3000 g for 10 mins at 4°C. Purified / CD71 enriched reticulocytes were mixed with equal number of erythrocytes (from the same blood sample, using material pelleted under the Percoll cushion) and prestained with Thiazole Orange (10 mg/ml stock solution, 1 μl from the stock added in 10ml of 0.2%BSA buffer). From this mix 0.5 million cells were added to the washed protein coated beads per well of 100 μl reaction volume and incubated for 45 mins at 4°C. The plate was centrifuged at 250 g for 10 mins and the supernatant containing unbound protein was removed carefully. The cell pellet was then resuspended gently with 0.2% BSA buffer and was centrifuged for 10 mins at 250 g, this washing step was repeated three times. The cells were finally resuspended in 200 μl 0.2% BSA and analysed using flow cytometry (BD Fortessa). Data were acquired using the FACS Diva software and post processing was performed using FlowJo (ver 10.2). PvRBP2b_169-813_ and Pfs25_23-193_ proteins were used as positive and negative controls respectively. Nile red and Thiazole Orange emissions were detected using 561 and 488 nm lasers respectively. The voltage settings for the respective lasers were FSC-367, SSC-293, 640/670-500, 488-433 and 561/582-480. The gating strategy has been shown in **Supplementary Figure 6**. The NR positive binding populations for both retics and erythrocytes has been calculated in terms of percentage with respect to the total retics (TO positive population) and total erythrocytes respectively.

### Structural Methods

#### Protein cloning, expression, and purification

All protein structural work was conducted prior to the availability of AlphaFold2. To identify constructs best suited for structural analysis, secondary structure composition for PVP01_0000100 was predicted using NetSurf 3.0 [61] identifying potential domain boundaries. This identified a C-terminal domain (CTD, D459-693L) of PVP01_0000100 possessing high helical content that was selected for expression and crystallisation studies. The PVP01_0000100_CTD was PCR amplified from the plasmid containing the PVP01_0000100 full length (FL, K62-693L) construct and cloned into the pHLSec vector with AgeI/KpnI restriction sites. The construct was expressed in 200 ml of HEK293F cells (Thermo) and purified with Ni-NTA (HiTrap, Cytiva) followed by buffer exchange in size exclusion buffer (20 mM Tris,150 mM NaCl, pH 7.4) for three times in a concentrator, MWCO 10 kDa (Milipore) prior to size exclusion (Superdex 75 16/600, analytical Hi load column, Cytiva). The protein was eluted and collected fractions were pooled together and concentrated using a centrifugal concentrator, 10 kDa MWCO (Millipore, Fischer scientific) to 10 mg/ml for crystallisation trials.

#### Crystallization and structure determination

Initial crystallisation trials were performed in 96-well plates using sitting drops composed of 200 nL protein plus 200 nL reservoir solution Initial crystals were identified in broad matrix screens (PACT PREMIER, Molecular Dimensions) and diffraction-quality crystals of Trag25_CTD grew in 0.1 M SPG, pH 4.0, 25% (w/v) PEG1500. The crystals were cryo-protected with 20% (v/v) glycerol added as supplement to the reservoir solution prior to flash-freezing in liquid nitrogen. Data were collected at the Diamond Light Source I04 beamline. Data processing and scaling was performed using Xia2 DIALS [62]. Phaser [63] was used for molecular replacement using a model generated from RoseTTAFold [64]. ArpWarp [65] was used to build an initial model of PVP01_0000100_CTD, followed by manual building in Coot [66]. Structure refinement was carried out with Coot [66], ISOLDE [67] and phenix.refine [68]. Data collection and refinement statistics are shown in **Supplementary Table 3**. Illustrations were generated using The PyMOL Molecular Graphics System, Version 2.0 Schrödinger, LLC. The surface charge analysis for the structure of PVP01_0000100_CTD was carried out with Adaptive Poisson -Boltzmann Solver (APBS) programme (server.poissonboltzmann.org) [69]. The results were analysed in UCSF chimera [70]. The atomic coordinates and structure factors have been deposited in the Protein Data Bank, www.pdb.org under accession code 8ARL.

### Lipid Dot blot

The membrane lipid strips (Echelon P-6002) were blocked with 3% fatty acid free BSA (Sigma, A4612) overnight at 4°C degree. The blocked membrane was incubated with 1 µg of PVP01_0000100_CTD, PVP01_0000100_FL and Pfs25 (negative control) for 2 h at RT. The proteins were incubated overnight followed by three washes with PBST. The strips were then incubated with anti-His-rabbit HRP 1:500 (Proteintech, Ptglab, USA). Detection was performed with an ECL reagent (GE Healthcare, USA).

### Liposome binding assay

Phosphotidylcholine (PC) was purchased from Merck (#840051C), Sulfatide was purchased from Avanti® (#131305P) and 1,2-dimyristoyl-sn-glycero-3-phosphoethanolamine-N- (lissamine rhodamine B sulfonyl) (Rhod-PE) was purchased from Avanti® (#810157P). Lipids were dissolved/suspended in chloroform and the final desired mixtures added to a glass vial (48% PC:50% sulfatide: 2% Rhod-PE and 98% PC: 2% Rhod-PE) before the chloroform was evaporated under a nitrogen stream. To all liposomes prepared, 2% Rhod-PE was incorporated to aid with liposome visualisation. The resultant lipid film was then hydrated using 50 mM HEPES (pH 7.4) and 150 mM NaCl to form multilamellar liposomes. A final liposome concentration of 0.8 mM in 50 µL was incubated with 1 µM protein for 30 min at room temperature with rotation. The liposomes were then pelleted via centrifugation at 20,000 g and the supernatant removed. The pellet was washed twice before resuspension in 10 µL of 50 mM HEPES (pH 7.4) and 150 mM NaCl and 10 µL of 2X loading dye and then ran on a NuPAGE™ 4-12% Bis-Tris Gel (Invitrogen, #NP0335) and stained with InstantBlue® (Abcam, #ISB1L).

### *In vitro* culture of *P. knowlesi*

Cultures of *P. knowlesi* were maintained as described previously [36]. Briefly, human O^+^ erythrocytes were used for the parasite culture and were purchased from NHSBT. The use of human cells for this work was approved by the NHS Cambridge South Research Ethics Committee (REC reference 20/EE/0100) and by the University of Cambridge Human Biology Research Ethics Committee (HBREC.2019.40). Parasites were grown in RPMI 1640 medium supplemented with Albumax (Thermo Fisher Scientific), L-Glutamine (Thermo Fisher Scientific), 10% Horse serum (Thermo Fisher Scientific), and Gentamicin (Thermo Fisher Scientific). The cultures were kept at 37°C with 2% haematocrit and kept in an incubator containing gas mixture of 3% CO_2_, 1% O_2_ and 96% Nitrogen. The cultures were monitored thrice a week by counting parasitaemia using light microscopy with media change or splitting as appropriate.

### Genetic modification of *P. knowlesi* parasites

#### PVP01_0000100 knockout construct design and assembly

The constructs, guide RNAs and primers were designed in Benchling. The 800 bp fragments immediately 5′ and 3′ upstream of PVP01_0000100 were amplified using *P. knowlesi* genomic DNA obtained from Pk infected erythrocytes using a DNA blood kit (Qiagen) according to the manufacturer’s protocol. The eGFP gene was amplified from a plasmid which is used in transfection as positive control (gift from Rob Moons lab). The 5′ and 3′ homology regions (88 and 85 ng/μl), eGFP gene (40 ng/μl) and pUC19 EcoR1/HindIII digested vector (70 ng/μl) were then added to the Gibson assembly mix at a 1:1:1 ratio and incubated for one hour at 50°C.

#### Cas9 vector and gRNA assembly

The cloning vector pKU6Cas9ccdB (1 μg) was digested and dephosphorylated using Fast Digest *Sap*I (*Lgu*I) (Thermo, FD1934) and Alkaline phosphatase (Thermo, EF0654) and ran on a 1% agarose gel followed by purification using agarose gel purification kit (Macherey and Nagel). The Guide RNAs were synthesised from Sigma and reconstituted by mixing 10 μM of the forward and reverse strands for each guide with 1 μl of 10X ligation buffer (NEB), 0.5 μl T4 polynucleotide kinase (NEB) and 6.5 μl nuclease free PCR water. Annealing was carried out by incubating at 37°C for 30 mins, then increasing to 95°C for 5 mins before cooling at 25°C at a ramp speed of 0.1°C/sec. Annealed primers were then diluted to 1 μl in 200 μl and ligated to the digested and dephosphorylated pKU6Cas9ccdB (50 ng/μl). Two gRNAs were constructed for PVP01_0000100 KO. The gene knockout vector was kindly provided by Rob Moon. The details of the reaction mixtures and primers for repair template and gRNA vector assemblies were provided in the **Supplementary Table 4**.

The assembled pUC19 Repair template and gRNA/Cas9 vector were transformed in *E. coli* Ultracompetent cells (XL-10 gold, Agilent) and plated onto Amp^+^ LB plates. Colonies positive for the insertion were sequence verified (Genewiz) before preparation using a midiprep purification kit (Macherey and Nagel) for transfection.

#### *P. knowlesi* synchronization, schizont enrichment and transfection

Synchronization was performed by enriching late-stage parasites using Histodenz (Sigma Aldrich) as described [71]. Briefly, the parasites were centrifuged at 1500 g, and the pellet was resuspended in 5 ml complete media and then layered on top of 5 ml of 55% Histodenz (prepared by adding 2.25 ml of complete medium + 2.75 ml of 100% Histodenz) in a 15 ml tube (Greiner). The mixture was then centrifuged for 10 mins at room temperature, 1500 g, acceleration 3 and brake 1, resulting in late-stage parasites becoming enriched at the interface. For immunofluorescence assays and transfections, this was repeated over three consecutive cell cycles to create tightly synchronized parasites, with schizont samples from a fourth cycle of Histodenz purification used for transfection. For each transfection, 20-25 μl of pure late schizonts were mixed with 90 μl of P3 solution in a cuvette (Lonza) containing 20 μg of repair template and 10 μg each of two guide vectors (total 40 µg). Transfections were carried out using program FP158 (Amaxa Nucleofector, Lonza), and the contents were immediately transferred into a 1.5 ml sterile Eppendorf tube containing 500 μl of prewarmed complete culture media and 150 μl of uninfected erythrocytes (50% Hct). The transfection mix was incubated at 37°C while shaking at 800 rpm in a thermomixer for 30 mins, before being transferred into a 6 well plate at 37°C for one parasite life cycle. After 24 hours, the parasites were subjected to 100 nM pyrimethamine (Santa Cruz Biotechnology Inc) selection media. The parasites were kept under selection for 7 days with drug media changed every two days. On day 7, the drug media was replaced with complete media. The smears of the parasites were checked for their recovery and parasitaemia around 1%. The parasite cultures were collected for gDNA extraction using a DNA Blood kit (Qiagen) for subsequent genotyping PCR to confirm editing. To clone out the WT population, limiting dilution and plaque cloning in flat-bottomed 96-well plates were performed. Wells containing single plaques were identified using an EVOS microscope (4X objective, transmitted light), expanded, and DNA isolated and genotyped as described above.

### Growth rate assays for genetically modified lines

Wild type and genetically modified *P. knowlsei* strains were synchronised as described above. Tightly synchronised ring parasites for both the strains were maintained in 6-well plates in normal complete RPMI media. Every alternate day, a 100 µL aliquot was collected, centrifuged and thickly smeared on the slide, before being fixed with methanol, air dried and stained with Giemsa for no longer than 10 mins. Cells were counted manually under the microscope with 100X oil immersion objective. A total of ∼5000 cells were counted for each slide for each line. The parasitaemia were determined and remaining culture was cut down to 1% parasitaemia and grown again. Data were collected this way for 14 days with 7 time points collected. The assay was done twice with technical replicates and analysed by plotting cumulative parasitaemia over time.

### Reticulocyte invasion assays

A 20 ml culture of *P. knowlesi* were synchronised as described above and schizonts were purified and washed in complete media. A 0.2 μl aliquot of schizonts from PkWT and PKNH_1300500 KO (∼90% pure) were added separately to two different tubes containing 4 μl (100% Hct) of enriched reticulocytes purified using CD71^+^ coated magnetic beads as described above. The final parasitaemia was ∼4.5% and the haematocrit was adjusted to 2% by adding 196 µl complete media (total volume 200 µl). A 50 μl volume of culture was added to each well and a set of 3 wells were set up for the PkWT and PKNH_1300500 KO line. The plate was incubated at 37°C for 12 hours to ensure at least one round of invasion. The next day, the cells were fixed in 100 μl of 2% PFA, 0.008% GA in PBS for 30 mins at 4°C followed by two times wash with PBS at 450 g, 3 mins each. The cells were then permeabilised for 10 mins at RT with 0.3% Triton X-100 in PBS followed by three washes with PBS. The cells were treated with 10 μg/ml Ribonuclease A in PBS per 100 μl (per well) for 1 h at 37°C followed by three washes with PBS. Finally, the cells were stained with SyBr green (1:5000) in PBS at 37°C for 30 mins followed by one wash with PBS at 450 g for 3 mins and finally resuspend in 100 μl PBS for analysis using flow cytometry (BD Fortessa).

### *P. vivax* ex vivo culture and immunofluorescence slide preparation

A cryopreserved Brazilian *P. vivax* isolate was thawed as reported previously [72]. The samples were then enriched on 1.080 g/ml KCl high Percoll gradients. Briefly, 2 ml of parasitised red blood cells resuspended using Iscove’s modified Dulbecco’s medium (IMDM) were layered on 3 ml 1.080 g/mL KCl high Percoll gradient and spun for 15 mins at 1200 *g* with slow acceleration and no break. The interface post Percoll-gradient spin was removed, washed in incomplete IMDM, and subjected to *in vitro* culture conditions with 1% hematocrit in IMDM (Gibco) containing 10% AB^+^ heat-inactivated sera and 50 μg/ml gentamicin at 37 °C in 5% CO_2,_ 1% O_2_, and 94% N_2_ [73]. For immunofluorescence assays, slide cytospins were made after 42-44 h of maturation, fixed in ice-cold methanol for 10 mins, air-dried and stored at -20°C.

### Immunofluorescence assays

To generate *P. knowlesi* slides for immunofluorescence, parasites were synchronised as described above and schizonts purified by density gradient centrifugation using a Histodenz (Sigma) cushion. They were then incubated with the Protein Kinase G inhibitor Compound 2 (1 µM) to prevent egress [74], resulting in the majority of parasites arresting as extreme late stage mature schizonts. After an hour of this treatment, the media was replaced with normal media for half an hour and finally washed with PBS and fixed for 30 min in 4% paraformaldehyde/0.0075% glutaraldehyde in PBS at room temperature. Fixing buffer was removed by PBS washing, and a schizont smear prepared and fixed on a glass slide. The smear was surrounded by hydrophobic ink from a PAP pen (Daido Sangyo, Japan) to allow low volume staining and washing. For permeabilization, the schizonts were incubated in permeabilization buffer (0.1% Triton X100 in PBS) for 30 mins at room temperature. To prevent nonspecific antibody interaction, the cells were blocked overnight at 4°C using a blocking buffer (3% BSA in PBS or 10% Goat Sera in PBS). For colocalisation studies (both for Pk and Pv), purified rabbit IgGs (described in the IgG purification section) against the selected TRAgs were used (in 1:750 dilution) along with rat PkMSP_1-19_ [18] With these primary antibodies, goat anti rabbit Alexa Fluor 633 (Thermo scientific) and goat anti rat Alexa Fluor 488 (Thermo Scientific) antibodies were used as secondary antibodies respectively in 1:500 dilutions. Primary antibodies were incubated with the samples sequentially for 1 hr each at room temperature. All the secondary antibodies and counterstain Hoechst 33342 (Thermo Fisher Scientific) together were incubated with the sample for 45 mins at room temperature in the dark. In between each incubation, washing was performed using PBS buffer with 0.05% Tween20 (3 times, 10 mins each). All primary and secondary antibodies were prepared in blocking buffer for incubation. After a final wash, the excess washing buffer was removed completely from the smear and the cells were left to mount with a mounting medium (Prolong gold antifade agent, Thermo Fisher) under a coverslip to cure overnight in the dark. The slides were either stored or visualised using a confocal microscope (LSM880 with airyscan setting). The captured images were processed using Zen Blue (2.0) software. The colocalization of PkMSP_1-19_ and the TRAGs antibodies were measured by determining the Pearson correlation coefficient (r) using additional JACoP in ImageJ.

### Flickering spectrometry of reticulocytes coated with proteins

50 µl of CD71+ reticulocytes were used as control and incubated at 4°C for 45 mins in PBS + 0.2% BSA with 100 µg/ml Pfs25 (negative control), and 100 µg/ml PVP01_0000100, respectively. Samples were washed once and diluted in PBS supplemented with 0.2% BSA at 0.01% Hct and transferred in a Secure Seal Hybridisation Chamber (Sigma-Aldrich, UK) attached to a glass cover slip; the same was kept at 37°C temperature through a heated collar. Cell membrane thermal fluctuations were recorded at a high frame rate (514 frames per s) and short exposure time (0.8 ms) for 20 s with a 60X Plan Apo VC 1.40 NA oil objective using a Nikon Eclipse Ti-E inverted microscope (Nikon, Japan). Videos were acquired in bright-field with a red filter using a CMOS camera (model GS3-U3-23S6M-C, Point Grey Research/FLIR Integrated Imaging Solutions (Machine Vision), Ri Inc., Canada). The equatorial contour for each frame was tracked by an in-house Matlab program as previously described [41, 75].

The radial component of each contour is then decomposed into its Fourier modes, and the average power spectrum for each cell is:

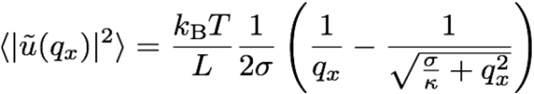

*u* is the amplitude of fluctuations, *q*_*x*_ is the mode, brackets denote averaging across all contours for a cell, *k*_*B*_ is the Boltzmann constant, *T* is the temperature, and *L* is the mean circumference of the cell contour. *σ* and *κ* are the tension and bending modulus, respectively, for which the same program fits the average power spectrum for mode numbers between 5 and 20, as done in [76]. Higher modes are dominated by noise, whereas lower modes are dominated by the shape of the cell.

### Cell membrane wrapping energy during invasion

The wrapping energy was calculated by considering changes to the free energy of the erythrocyte membrane due to a localised deformation and depends on its tension and bending modulus. We consider the deformation to be localised to a disc of radius R which is deformed by a merozoite to a half-sphere of the same radius. Assuming a very simple hemi-spherical geometry, the sum of these two contributions is given by

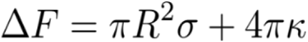

where *R* = 1 μm is the radius of the merozoite [77, 78], *σ* is the tension, and *κ* is the bending modulus. The above equation is derived in [41].

## Data Availability

The authors declare that the data supporting the findings of this study are available within this Manuscript, Supplementary Information and Methods. All AF2 models and associated statistics will be deposited in the University of Cambridge Data Repository. The structure factors and atomic coordinates for PVP01_0000100 C terminal domain are deposited in the PDB under the accession code 8ARL.

## Acknowledgements

We would like to thank Prof. Gavin J. Wright for his help during the construct designing for expression vectors. Dr. Reiner Schulte, Gabriela Grondys-Kotarba and Chiara Cossetti of the CIMR Flow Cytometry facility for providing required training and assistance during the flow cytometry experiments. We also like to convey our thanks to Matthew Gratian and Mark Bowen for providing training and data acquisition in LSM880 confocal microscope. We are grateful to Alison Kemp for her guidance to use the CRISPR Cas9 system. Ellen Knuepfer for the kind donation of Rat anti-PkMSP1-19 serum. We thank Stephen Graham for help with crystallographic model building. This work was funded by the National Institutes of Health (R01AI137154) and the Wellcome Trust (222323/Z/21/Z). J.E.D and S.J.M. are supported by a Wellcome Trust Senior Research Fellowship (219447/Z/19/Z) awarded to J.E.D. As this research was funded in part by the Wellcome Trust, for the purpose of open access, the author has applied a CC BY public copyright license to any Author Accepted Manuscript version arising from this submission.

## Author contributions

J.C.R, J.E.D, M.D. and P.K. conceived and designed the research; P.K., D.N., S.M., V.I., performed the experiments, P.K., D.N., S.M., V.I, M.D., J.E.D & J.C.R analysed the data; P.K. and J.E.D. performed structural refinement and analysis, M.D., S.D. and U.K. provided the Pv isolates for the IFA studies. P.K. J.C.R., D.N., J.E.D., S.M wrote the first draft of the manuscript and all authors provided input.

## References

1. Battle, K.E., et al., Mapping the global endemicity and clinical burden of Plasmodium vivax, 2000-17: a spatial and temporal modelling study. Lancet, 2019. 394(10195): p. 332–343.

2. Global technical strategy for malaria 2016-2030. 2015.

3. Crosnier, C., et al., Basigin is a receptor essential for erythrocyte invasion by Plasmodium falciparum. Nature, 2011. 480(7378): p. 534–7.

4. Douglas, A.D., et al., A PfRH5-based vaccine is efficacious against heterologous strain blood-stage Plasmodium falciparum infection in aotus monkeys. Cell Host Microbe, 2015. 17(1): p. 130–9.

5. Minassian, A.M., et al., Reduced blood-stage malaria growth and immune correlates in humans following RH5 vaccination. Med (N Y), 2021. 2(6): p. 701–719 e19.

6. Singh, K., et al., Malaria vaccine candidate based on Duffy-binding protein elicits strain transcending functional antibodies in a Phase I trial. NPJ Vaccines, 2018. 3: p. 48.

7. Lo, E., et al., Frequent expansion of Plasmodium vivax Duffy Binding Protein in Ethiopia and its epidemiological significance. PLoS Negl Trop Dis, 2019. 13(9): p. e0007222.

8. Gruszczyk, J., et al., Transferrin receptor 1 is a reticulocyte-specific receptor for Plasmodium vivax. Science, 2018. 359(6371): p. 48–55.

9. Cowman, A.F. and B.S. Crabb, Invasion of red blood cells by malaria parasites. Cell, 2006. 124(4): p. 755–66.

10. Burns, J.M., et al., An unusual tryptophan-rich domain characterizes two secreted antigens of Plasmodium yoelii-infected erythrocytes. Mol Biochem Parasitol, 2000. 110(1): p. 11–21.

11. Oguariri, R.M., et al., High prevalence of human antibodies to recombinant Duffy binding-like alpha domains of the Plasmodium falciparum-infected erythrocyte membrane protein 1 in semi-immune adults compared to that in nonimmune children. Infect Immun, 2001. 69(12): p. 7603–9.

12. Zeeshan, M., et al., Host-parasite interaction: selective Pv-fam-a family proteins of Plasmodium vivax bind to a restricted number of human erythrocyte receptors. J Infect Dis, 2015. 211(7): p. 1111–20.

13. Alam, M.S., et al., Interaction of Plasmodium vivax Tryptophan-rich Antigen PvTRAg38 with Band 3 on Human Erythrocyte Surface Facilitates Parasite Growth. J Biol Chem, 2015. 290(33): p. 20257–72.

14. Tyagi, K., et al., Plasmodium vivax Tryptophan Rich Antigen PvTRAg36.6 Interacts with PvETRAMP and PvTRAg56.6 Interacts with PvMSP7 during Erythrocytic Stages of the Parasite. PLoS One, 2016. 11(3): p. e0151065.

15. Gunalan, K., et al., Transcriptome profiling of Plasmodium vivax in Saimiri monkeys identifies potential ligands for invasion. Proc Natl Acad Sci U S A, 2019. 116(14): p. 7053–7061.

16. Wang, B., et al., Immunoprofiling of the Tryptophan-Rich Antigen Family in Plasmodium vivax. Infection and Immunity, 2015. 83(8): p. 3083–3095.

17. Hostetler, J.B., et al., A Library of Plasmodium vivax Recombinant Merozoite Proteins Reveals New Vaccine Candidates and Protein-Protein Interactions. PLoS Negl Trop Dis, 2015. 9(12): p. e0004264.

18. Ndegwa, D.N., et al., Using Plasmodium knowlesi as a model for screening Plasmodium vivax blood-stage malaria vaccine targets reveals new candidates. PLoS Pathog, 2021. 17(7): p. e1008864.

19. Ntumngia, F.B., N. Bahamontes-Rosa, and J.F. Kun, Genes coding for tryptophan-rich proteins are transcribed throughout the asexual cycle of Plasmodium falciparum. Parasitol Res, 2005. 96(6): p. 347–53.

20. Bahl, A., et al., PlasmoDB: the Plasmodium genome resource. A database integrating experimental and computational data. Nucleic Acids Res, 2003. 31(1): p. 212–5.

21. Kall, L., A. Krogh, and E.L. Sonnhammer, Advantages of combined transmembrane topology and signal peptide prediction--the Phobius web server. Nucleic Acids Res, 2007. 35(Web Server issue): p. W429–32.

22. Hiller, K., et al., PrediSi: prediction of signal peptides and their cleavage positions. Nucleic Acids Res, 2004. 32(Web Server issue): p. W375–9.

23. Crosnier, C., et al., A library of functional recombinant cell-surface and secreted P. falciparum merozoite proteins. Mol Cell Proteomics, 2013. 12(12): p. 3976–86.

24. Ntumngia, F.B., et al., Characterisation of a tryptophan-rich Plasmodium falciparum antigen associated with merozoites. Mol Biochem Parasitol, 2004. 137(2): p. 349–53.

25. Barr, P.J., et al., Recombinant Pfs25 protein of Plasmodium falciparum elicits malaria transmission-blocking immunity in experimental animals. J Exp Med, 1991. 174(5): p. 1203–8.

26. Kaslow, D.C., et al., Comparison of the primary structure of the 25 kDa ookinete surface antigens of Plasmodium falciparum and Plasmodium gallinaceum reveal six conserved regions. Mol Biochem Parasitol, 1989. 33(3): p. 283–7.

27. Vermeulen, A.N., et al., Sequential expression of antigens on sexual stages of Plasmodium falciparum accessible to transmission-blocking antibodies in the mosquito. J Exp Med, 1985. 162(5): p. 1460–76.

28. Jumper, J., et al., Highly accurate protein structure prediction with AlphaFold. Nature, 2021. 596(7873): p. 583–589.

29. Holm, L., Using Dali for Protein Structure Comparison. Methods Mol Biol, 2020. 2112: p. 29–42.

30. Peter, B.J., et al., BAR domains as sensors of membrane curvature: the amphiphysin BAR structure. Science, 2004. 303(5657): p. 495–9.

31. Gallop, J.L., et al., Mechanism of endophilin N-BAR domain-mediated membrane curvature. EMBO J, 2006. 25(12): p. 2898–910.

32. Masuda, M., et al., Endophilin BAR domain drives membrane curvature by two newly identified structure-based mechanisms. EMBO J, 2006. 25(12): p. 2889–97.

33. Zimmerberg, J. and S. McLaughlin, Membrane curvature: how BAR domains bend bilayers. Curr Biol, 2004. 14(6): p. R250–2.

34. Roberts, D.D., et al., Laminin binds specifically to sulfated glycolipids. Proc Natl Acad Sci U S A, 1985. 82(5): p. 1306–10.

35. Zhou, Z., et al., Erythrocyte membrane sulfatide plays a crucial role in the adhesion of sickle erythrocytes to endothelium. Thromb Haemost, 2011. 105(6): p. 1046–52.

36. Moon, R.W., et al., Adaptation of the genetically tractable malaria pathogen Plasmodium knowlesi to continuous culture in human erythrocytes. Proc Natl Acad Sci U S A, 2013. 110(2): p. 531–6.

37. Malleret, B., et al., Significant biochemical, biophysical and metabolic diversity in circulating human cord blood reticulocytes. PLoS One, 2013. 8(10): p. e76062.

38. Amir, A., et al., Invasion characteristics of a Plasmodium knowlesi line newly isolated from a human. Sci Rep, 2016. 6: p. 24623.

39. Yoon, Y.Z., et al., Flickering analysis of erythrocyte mechanical properties: dependence on oxygenation level, cell shape, and hydration level. Biophys J, 2009. 97(6): p. 1606–15.

40. Kariuki, S.N., et al., Red blood cell tension protects against severe malaria in the Dantu blood group. Nature, 2020. 585(7826): p. 579–583.

41. Introini, V., et al., The erythrocyte membrane properties of beta thalassaemia heterozygotes and their consequences for Plasmodium falciparum invasion. Sci Rep, 2022. 12(1): p. 8934.

42. Gardner, M.J., et al., Genome sequence of the human malaria parasite Plasmodium falciparum. Nature, 2002. 419(6906): p. 498–511.

43. Auburn, S., et al., A new Plasmodium vivax reference sequence with improved assembly of the subtelomeres reveals an abundance of pir genes. Wellcome Open Res, 2016. 1: p. 4.

44. Frost, A., et al., Structural basis of membrane invagination by F-BAR domains. Cell, 2008. 132(5): p. 807–17.

45. Dawson, J.C., J.A. Legg, and L.M. Machesky, Bar domain proteins: a role in tubulation, scission and actin assembly in clathrin-mediated endocytosis. Trends Cell Biol, 2006. 16(10): p. 493–8.

46. Lyon, A.M., et al., Full-length Galpha(q)-phospholipase C-beta3 structure reveals interfaces of the C-terminal coiled-coil domain. Nat Struct Mol Biol, 2013. 20(3): p. 355–62.

47. Cerami, C., F. Kwakye-Berko, and V. Nussenzweig, Binding of malarial circumsporozoite protein to sulfatides [Gal(3-SO4)beta 1-Cer] and cholesterol-3-sulfate and its dependence on disulfide bond formation between cysteines in region II. Mol Biochem Parasitol, 1992. 54(1): p. 1–12.

48. Pancake, S.J., et al., Malaria sporozoites and circumsporozoite proteins bind specifically to sulfated glycoconjugates. J Cell Biol, 1992. 117(6): p. 1351–7.

49. Namvar, A., et al., Surface area-to-volume ratio, not cellular viscoelasticity, is the major determinant of red blood cell traversal through small channels. Cell Microbiol, 2021. 23(1): p. e13270.

50. Burns, J.M., Jr., E.K. Adeeku, and P.D. Dunn, Protective immunization with a novel membrane protein of Plasmodium yoelii-infected erythrocytes. Infect Immun, 1999. 67(2): p. 675–80.

51. Julie-Anne Gabelich, J.G., Florian Kirscht, Oliver Popp, Joachim M Matz, Gunnar Dittmar, Melanie Rug, Alyssa Ingmundson, A member of the tryptophan-rich protein family is required for efficient sequestration of Plasmodium berghei schizonts. bioRxiv, 2022.

52. Bahl, A., et al., PlasmoDB: the Plasmodium genome resource. An integrated database providing tools for accessing, analyzing and mapping expression and sequence data (both finished and unfinished). Nucleic Acids Res, 2002. 30(1): p. 87–90.

53. Sievers, F., et al., Fast, scalable generation of high-quality protein multiple sequence alignments using Clustal Omega. Mol Syst Biol, 2011. 7: p. 539.

54. Dereeper, A., et al., Phylogeny.fr: robust phylogenetic analysis for the non-specialist. Nucleic Acids Res, 2008. 36(Web Server issue): p. W465–9.

55. Felsenstein, J., Evolutionary trees from DNA sequences: a maximum likelihood approach. J Mol Evol, 1981. 17(6): p. 368–76.

56. Felsenstein, J., Confidence Limits on Phylogenies: An Approach Using the Bootstrap. Evolution, 1985. 39(4): p. 783–791.

57. Petersen, T.N., et al., SignalP 4.0: discriminating signal peptides from transmembrane regions. Nat Methods, 2011. 8(10): p. 785–6.

58. Sonnhammer, E.L., G. von Heijne, and A. Krogh, A hidden Markov model for predicting transmembrane helices in protein sequences. Proc Int Conf Intell Syst Mol Biol, 1998. 6: p. 175–82.

59. Crosnier, C., N. Staudt, and G.J. Wright, A rapid and scalable method for selecting recombinant mouse monoclonal antibodies. BMC Biol, 2010. 8: p. 76.

60. Sorette, M.P., K. Shiffer, and M.R. Clark, Improved isolation of normal human reticulocytes via exploitation of chloride-dependent potassium transport. Blood, 1992. 80(1): p. 249–54.

61. Hoie, M.H., et al., NetSurfP-3.0: accurate and fast prediction of protein structural features by protein language models and deep learning. Nucleic Acids Res, 2022.

62. Kabsch, W., Xds. Acta Crystallogr D Biol Crystallogr, 2010. 66(Pt 2): p. 125–32.

63. McCoy, A.J., et al., Phaser crystallographic software. J Appl Crystallogr, 2007. 40(Pt 4): p. 658–674.

64. Baek, M., et al., Accurate prediction of protein structures and interactions using a three-track neural network. Science, 2021. 373(6557): p. 871–876.

65. Langer, G., et al., Automated macromolecular model building for X-ray crystallography using ARP/wARP version 7. Nat Protoc, 2008. 3(7): p. 1171–9.

66. Emsley, P., et al., Features and development of Coot. Acta Crystallogr D Biol Crystallogr, 2010. 66(Pt 4): p. 486–501.

67. Croll, T.I., ISOLDE: a physically realistic environment for model building into low-resolution electron-density maps. Acta Crystallogr D Struct Biol, 2018. 74(Pt 6): p. 519–530.

68. Afonine, P.V., et al., Towards automated crystallographic structure refinement with phenix.refine. Acta Crystallogr D Biol Crystallogr, 2012. 68(Pt 4): p. 352–67.

69. Jurrus, E., et al., Improvements to the APBS biomolecular solvation software suite. Protein Sci, 2018. 27(1): p. 112–128.

70. Pettersen, E.F., et al., UCSF Chimera--a visualization system for exploratory research and analysis. J Comput Chem, 2004. 25(13): p. 1605–12.

71. Moon, R.W., et al., Normocyte-binding protein required for human erythrocyte invasion by the zoonotic malaria parasite Plasmodium knowlesi. Proc Natl Acad Sci U S A, 2016. 113(26): p. 7231–6.

72. de Oliveira, T.C., et al., Genome-wide diversity and differentiation in New World populations of the human malaria parasite Plasmodium vivax. PLoS Negl Trop Dis, 2017. 11(7): p. e0005824.

73. Rangel, G.W., et al., Enhanced Ex Vivo Plasmodium vivax Intraerythrocytic Enrichment and Maturation for Rapid and Sensitive Parasite Growth Assays. Antimicrob Agents Chemother, 2018. 62(4).

74. Collins, C.R., et al., Malaria parasite cGMP-dependent protein kinase regulates blood stage merozoite secretory organelle discharge and egress. PLoS Pathog, 2013. 9(5): p. e1003344.

75. Introini, V., et al., Biophysical Tools and Concepts Enable Understanding of Asexual Blood Stage Malaria. Front Cell Infect Microbiol, 2022. 12: p. 908241.

76. Koch, M., et al., Plasmodium falciparum erythrocyte-binding antigen 175 triggers a biophysical change in the red blood cell that facilitates invasion. Proc Natl Acad Sci U S A, 2017. 114(16): p. 4225–4230.

77. Dasgupta, S., et al., Membrane-wrapping contributions to malaria parasite invasion of the human erythrocyte. Biophys J, 2014. 107(1): p. 43–54.

78. Dasanna, A.K., et al., Effect of malaria parasite shape on its alignment at erythrocyte membrane. Elife, 2021. 10.

